# Testosterone use in female mice does not impair fertilizability of eggs: Implications for the fertility care of transgender males

**DOI:** 10.1101/2020.04.09.033803

**Authors:** C.B. Bartels, T.F. Uliasz, L. Lestz, L.M. Mehlmann

## Abstract

**STUDY QUESTION:** Does testosterone use in females affect reproductive potential, particularly with regard to the production of fertilizable gametes?

**SUMMARY ANSWER:** Testosterone cypionate injections given to post-pubertal female mice caused virilization and ovaries were smaller than control ovaries, but ovaries were still responsive to hormonal stimulation and produced fertilizable eggs when superovulated.

**WHAT IS KNOWN ALREADY:** Studies to examine the effects of testosterone on reproductive potential in transgender males are lacking. Recently, a model was developed that simulates many aspects of testosterone use in transgender males in order to look at reproductive effects of testosterone in female mice. This study found masculinizing effects on the mice but did not find significant deficits on the number of ovarian follicles; however, effects of testosterone use on ovarian stimulation and fertilizability of oocytes were not investigated.

**STUDY DESIGN, SIZE, DURATION:** A total of 66, 6-week-old Hsd:NSA(CF-1) female mice and 6 Hsd:ICR (CD-1) mice were used for this study. Mice were injected subcutaneously with 400 μg testosterone cypionate or sesame oil once a week for 6 weeks and were either sacrificed a week after the 6^th^ injection (active exposure group), or were sacrificed 6-7 weeks after the final testosterone injection (washout group).

**PARTICIPANTS/MATERIALS, SETTING, METHODS:** Both active exposure and washout groups were further subdivided into 3 groups: unstimulated, eCG-stimulated, or eCG/hCG-stimulated. eCG-stimulated mice were sacrificed 44-48 hrs after eCG injection. eCG/hCG-stimulated mice were injected with eCG, followed 48 hrs later with hCG. Mice were sacrificed ∼13-18 hrs after the hCG injection. Data collected included daily vaginal cytology, terminal hormone levels and ovary weights, ovarian histology, number of oocytes/eggs collected in each group, and cleavage to the 2-cell stage following in vitro fertilization.

**MAIN RESULTS AND THE ROLE OF CHANCE:** Testosterone cypionate-treated mice had testosterone levels elevated to the level of male mice and ceased cycling. Ovaries were significantly smaller in testosterone-treated mice, but they contained normal cohorts of follicles and responded to gonadotropin stimulation by ovulating similar numbers of eggs that fertilized and cleaved in vitro.

**LIMITATIONS, REASONS FOR CAUTION:** Our model treated female mice for only 6 weeks, whereas many transgender men use testosterone for many years before considering biological children. Importantly, a mouse system may not perfectly simulate human reproductive physiology.

**WIDER IMPLICATIONS OF THE FINDINGS:** The current standard of care for transgender men who desire biological children is to cease testosterone therapy prior to ovarian stimulation, but the necessity for stopping testosterone is not known. Our model demonstrates that it is possible for testosterone-suppressed ovaries to respond to gonadotropic stimulation by producing and ovulating fertilizable eggs, thereby obviating the need for testosterone cessation prior to ovarian stimulation. In time, these results may provide insights for future clinical trials of fertility treatment options for transgender men.

## Introduction

Transgender males are individuals who were assigned female at birth but identify as males. Many, but not all, transgender males opt to undergo gender-affirming treatment, which can consist of surgery and/or hormone therapy (HT) by long-term administration of testosterone (Quinn et al., 2017). HT improves gender dysphoria through testosterone-driven development and maintenance of desired male secondary sex characteristics; however, a potential adverse effect of testosterone exposure is a decrease in fertility.

A recent study by the Williams Institute estimated that about 1.4 million individuals identify as transgender in the United States (Flores, 2016), and there are reports that approximately half of transgender adults desire biological children (Moravek, 2019; Wierckx et al., 2012). The reproductive consequences of HT are still unclear, and both the World Professional Association for Transgender Health (WPATH) and the Endocrine Society recommend that all transgender males be counseled regarding options for fertility preservation before initiating testosterone therapy (Hembree et al., 2017; Meyer, 2009). Transgender males may not consider fertility preservation to be important at the start of testosterone therapy, which can be initiated as early as 14 years old. In addition, assisted reproductive technology centers have little experience in stimulation of peripubertal ovaries, nor in performing transvaginal oocyte harvest in children. Due to a variety of physiological and psychological barriers, ovarian stimulation and oocyte harvest is best avoided in children, if it can be safely postponed to adulthood. A recent study showed that only 2 of 72 (2.8%) young transgender individuals chose to utilize fertility preservation after counseling (Nahata et al., 2019), which reflects a priority for HT initiation to attain features of their affirmed gender while avoiding the delay, invasiveness, or costs of fertility preservation (Armuand et al., 2017; Insogna et al., 2020). The desire for biological children may arise later in life, after months or years of HT exposure.

The approach to fertility options in transgender males already taking HT therefore warrants more investigation. To date, studies to evaluate the impact of HT on reproductive potential for transgender males are lacking (ASRM Committee Opinion, 2015). The options for transgender males presenting for fertility preservation after HT is either surgical oophorectomy to collect ovarian tissue or surgical oocyte retrieval following ovarian stimulation (De Roo et al., 2016; Neblett and Hipp, 2019). Methods for maturation and fertilization of oocytes collected directly from isolated ovarian tissue without hormonal stimulation, while improving, are still considered to be experimental (Yang and Chian, 2018) and to date, there have been no studies to examine this method for fertility preservation in transgender males. Accordingly, if a transgender male presents for fertility treatment now or in the near future and plans to have the pregnancy carried by a cis-female partner or gestational carrier, the best option is in vitro fertilization (IVF) after ovarian stimulation and oocyte retrieval. Due to the unknown effects of high-level testosterone on ovarian response and oocyte quality, the current recommended practice before IVF is discontinuation of testosterone to allow the resumption of menses (Adeleye et al., 2019; Broughton and Omurtag, 2017; Leung et al., 2018). While this treatment regimen can be effective, HT cessation for the purpose of fertility treatment has been reported to cause significant psychological distress in the form of gender dysphoria attributed to the gender-incongruous effects of testosterone withdrawal, estrogen exposure, and menses (Armuand et al., 2017). These negative consequences could lead to treatment avoidance even when fertility is desired.

Ovarian tissue taken from HT-exposed transgender males has demonstrated changes including a thickened cortex, stromal hyperplasia, an increased number of atretic follicles, and increased cortical stiffness (De Roo et al., 2019; Ikeda et al., 2013). However, the ovarian tissue follicular pool is not diminished (De Roo et al., 2016; Van Den Broecke et al., 2001) Markers of ovarian reserve, including anti-Müllerian hormone and inhibin, are unchanged (Rodriguez-Wallberg et al., 2014), and successful pregnancies have been reported after testosterone use (Light et al., 2014). Case reports have been published of subjects successfully undergoing IVF after temporarily discontinuing testosterone therapy for 1-12 months, and healthy live births were reported (Adeleye et al., 2019; Broughton and Omurtag, 2017; Leung et al., 2018). These data suggest that the follicular pool and oocyte quality are preserved.

A primary mouse model for HT in transgender males was recently published and found that ovaries from testosterone-treated mice were generally normal, with the exception of some cyst-like late antral follicles (Kinnear et al., 2019). The fertility potential in terms of ovarian response to gonadotropins, oocyte integrity, or fertilizability was not examined. There is very limited information about the necessity for cessation of testosterone therapy prior to ovarian stimulation, and no mouse models have addressed this problem. The aim of the present study was to establish a mouse model in which reproductive potential could be evaluated following prolonged HT in female mice with and without a period of testosterone cessation.

## Materials and Methods

### Ethical approval

Animal studies were performed in accordance with the Guide for the Care and Use of Laboratory Animals (National Academy of Sciences 1996) and were approved by the Institutional Animal Care & Use Committee at UConn Health (protocol number 101977-0122).

### Media and Reagents

All chemicals were purchased from Millipore Sigma (St. Louis, MO, USA) unless otherwise indicated. Testosterone cypionate was from Steraloids (Newport, RI, USA) and was prepared as an 8 mg/ml solution in sesame oil. Equine chorionic gonadotropin (eCG) was from Calbiochem. The medium for oocyte collection was HEPES-buffered MEMα (Gibco 12000022, Thermo Fisher, Waltham, MA, USA) containing penicillin, streptomycin, and polyvinyl alcohol (PVA), and 10 μM milrinone to prevent spontaneous meiotic maturation (Mehlmann et al., 2019). For overnight oocyte maturation, oocytes were washed into bicarbonate-buffered MEMα (Mehlmann et al., 2019) containing 5% fetal bovine serum (Invitrogen, Carlsbad, CA, USA) without milrinone. For in vitro fertilization, cumulus masses were collected in human tubal fluid medium (HTF; Cook Medical Inc. IVF medium (#K-RVFE; Fisher) containing reduced glutathione. Sperm were capacitated in IVF medium (Mehlmann and Kline, 1994) containing 15 mg/ml Fraction V bovine serum albumin.

### Experimental design

Six-week-old female CF-1 (Envigo, Indianapolis, IN, USA) and >8-week-old male CD-1 mice (Envigo) were used for all experiments. Three female mice were housed per cage in a temperature and light-controlled room on a 14L:10D light cycle. Male mice were housed individually.

Six-week-old female mice were lightly sedated with isoflurane and injected weekly, subcutaneously, with 400 μg testosterone cypionate or vehicle using 27-gauge needles. In the first set of experiments, mice were sacrificed within 8 days after the 6^th^ testosterone injection, when testosterone levels were high. These mice are referred to as the “active exposure” group. In the second set of experiments, mice were sacrificed 6-7 weeks after the 6^th^ testosterone injection, when testosterone returned to basal levels. These mice are referred to as the “washout” group. Both groups were subdivided into 3 more groups: mice that were not stimulated with gonadotropins; mice that were stimulated with eCG only; and mice that were stimulated with eCG followed by hCG to induce ovulation.

For both the active exposure and washout groups, we performed vaginal smears to examine cyclicity. We analyzed the following: serum testosterone levels; serum estrogen levels; ovary weights and histology; oocyte number prior to and after priming with eCG or eCG + hCG; structure of the meiotic spindle; and egg fertilizability. Control mice were injected with sesame oil and were treated in parallel with the testosterone-injected groups. All mice were euthanized by isoflurane overdose followed by cervical dislocation.

### Vaginal cytology

For the active exposure group, daily vaginal smears were performed using standard methods (Goldman et al., 2007) starting in the 5^th^ week of testosterone treatments. For the washout group, daily vaginal smears were done starting one week after the final testosterone injection. Staging of the estrous cycle was determined by the presence and distribution of leukocytes, cornified epithelial cells, and nucleated epithelial cells. Proestrus was identified by nucleated epithelium, estrus was identified by large cornified epithelial cells, metestrus was identified by leukocytes and large cornified epithelial cells, and diestrus was identified by the predominance of leukocytes in the presence of nucleated and cornified cells (Gaytan et al., 2017). Clitoral size was visually assessed at the time of cytology.

### Blood collection and hormone analysis

After the mice were sacrificed, they were weighed and terminal blood was collected by cardiac puncture using a heparinized 18-gauge needle and syringe. Blood samples were kept on ice. Within 30 minutes of collection, samples were centrifuged at 4°C for 15 minutes (1000 x *g*) and supernatants were stored at −80°C. Diethyl ether extraction was performed as directed by Cayman Chemical, with extracted samples stored at −20°C in Cayman ELISA buffer. Testosterone and estradiol-17β analyses were performed using ELISA kits (Cayman Chemical, Ann Arbor, MI, USA) according to the manufacturer’s instructions. For comparison, we also collected blood from mature male mice that were used for in vitro fertilization (see below). These males had been acclimated to the lab for at least one week prior to the experiment, were housed individually, and were not exposed to females prior to blood collection.

### Ovarian histology

Ovaries were collected and most of the fat and oviducts were removed by dissection under a stereoscope and were then weighed. Ovaries were fixed in 10% formalin for 24-48 hours, washed into PBS, then were dehydrated in ethanol, embedded in paraffin, and 5 μm serial sections were cut and processed by the Histology Core at UConn Health. Sections were stained with hematoxylin and eosin.

Total numbers of antral follicles and corpora lutea were counted. Antral follicles were defined as being ∼250-320 μm in diameter with a clearly visible antral cavity and oocyte with two or more layers of granulosa cells. Preovulatory follicles were defined as being >320 μm in diameter. Each antral follicle was counted only when the oocyte was present and while scanning between adjacent sections to prevent duplicate counting. Corpora lutea were defined as discrete eosinophilic round structures. Corpora lutea were numbered as the sections were serially assessed through the entire ovary to prevent duplicate counting. Two of the investigators independently counted follicles.

### Oocyte collection and immunofluorescence staining

Ovaries from unstimulated and eCG-primed mice were weighed and one ovary from each mouse was fixed for histological analysis while the contralateral ovary was used for oocyte collection. The ovary for oocyte collection was placed in HEPES-buffered MEMα containing milrinone and punctured using a 30-gauge needle. Oocytes were collected with a mouth pipet and counted. For in vitro maturation, oocytes were washed into bicarbonate-buffered MEMα without milrinone and were incubated overnight at 37°C in a humidified incubator containing 5% CO_2_/95% air. In vitro maturation was confirmed by the disappearance of the nuclear envelope and the formation of first polar bodies using a stereoscope. Oocytes were fixed for 30-60 min at 37°C in 2% formaldehyde, 100 mM HEPES, 50 mM EGTA, 10 mM MgSO_4_, and 0.2% Triton X-100, then were permeabilized in PBS containing 0.1% Triton X-100, and blocked for at least 15 minutes in PBS containing 3% BSA and 0.01% Triton X-100. Oocytes were incubated overnight at 4°C in primary antibody against tubulin (YL1/2; Serotec Inc., Raleigh, NC) diluted to 10 μg/ml in blocking buffer. After washing in PBS-PVA, oocytes were incubated in Alexa488-conjugated secondary antibody for 1 hr at room temperature in the dark. Oocytes were washed in PBS-PVA containing 5 μM SYTOX Orange (ThermoFisher) to label chromosomes. Imaging for spindle integrity was performed using a Zeiss Pascal confocal microscope with a 40X, 1.2 NA water immersion objective (C-Apochromat; Carl Zeiss MicroImaging, Inc., Thornwood, NY, USA).

### In vitro fertilization

Female mice were superovulated with 5 IU eCG, followed 48 hrs later with 5 IU hCG. Approximately 13-15 hrs later, ovaries and oviducts were removed and cumulus masses obtained by puncturing the swollen ampullae. Cumulus masses were incubated in 200 μl drops of HTF containing reduced glutathione for ∼30 min prior to adding sperm. Sperm were collected from the epididymides of male mice by gently snipping with fine scissors into a 100 μl drop of capacitation medium and were capacitated for 1-2 hrs before adding 3-5 μl of the sperm suspension to the drops containing the eggs. The sperm and eggs were incubated together for 4 hrs, then were washed into 200 μl drops of HTF without glutathione. Fertilized eggs were incubated overnight in a humidified incubator containing 5% CO2/95% air. The next day, 2-cell embryos were counted.

### Statistical analysis

Statistical analyses were performed using Prism 6.0 software for Windows, GraphPad Software, La Jolla, California (www.graphpad.com). Specific statistical tests for each experiment are indicated in the figure legends. P<0.05 was considered to be significant.

## Results

### Testosterone cypionate elevates serum testosterone levels and induces virilization in female mice

In a recent study investigating the effects of testosterone on female mice, Kinnear et al. (2019) injected testosterone enanthate twice weekly to maintain elevated testosterone levels. In the current study, we injected a similar form of testosterone, testosterone cypionate (referred to hereafter as “T”), which is commonly used by transgender men to elevate T levels (Luthy et al., 2017; Moravek et al., 2020), weekly. T-injected mice showed signs of virilization, including distinct clitoromegaly and cessation of estrous cycles (Fig. 1A). All control-injected mice clearly cycled throughout the entire experiment, whereas all T-injected mice appeared to be in diestrus (Fig.1B), which is consistent with what was observed previously (Kinnear et al., 2019). One week after the 6^th^ T injection (referred to herein as the “active exposure” group), mice were sacrificed, trunk blood was collected, and T levels were measured. T-injected mice had significantly higher T levels than controls, and the amount of T was comparable to the levels in adult males (Fig. 2). In one set of experiments, we did not sacrifice females after the 6-week injection period; rather, the mice were kept for several weeks after cessation of injections (referred to herein as the “washout” group). T levels declined to baseline levels within 5 weeks following the last injection (Fig. 2) and these “washout mice” resumed cycling, as assessed by daily vaginal smears. Interestingly, clitoromegaly was no longer apparent in these washout mice 5 weeks after the last injection (Fig. 1A). The weights of the mice did not differ between T-treated and controls. T-treated and control mice in the active exposure group weighed 30.5 ± 0.8 g (SEM) vs 31 ±1 g, respectively, whereas in the washout group T-treated and control mice weighed 33.9 ± 1 and 33.7 ± 0.8 g, respectively.

**Figure 1.**
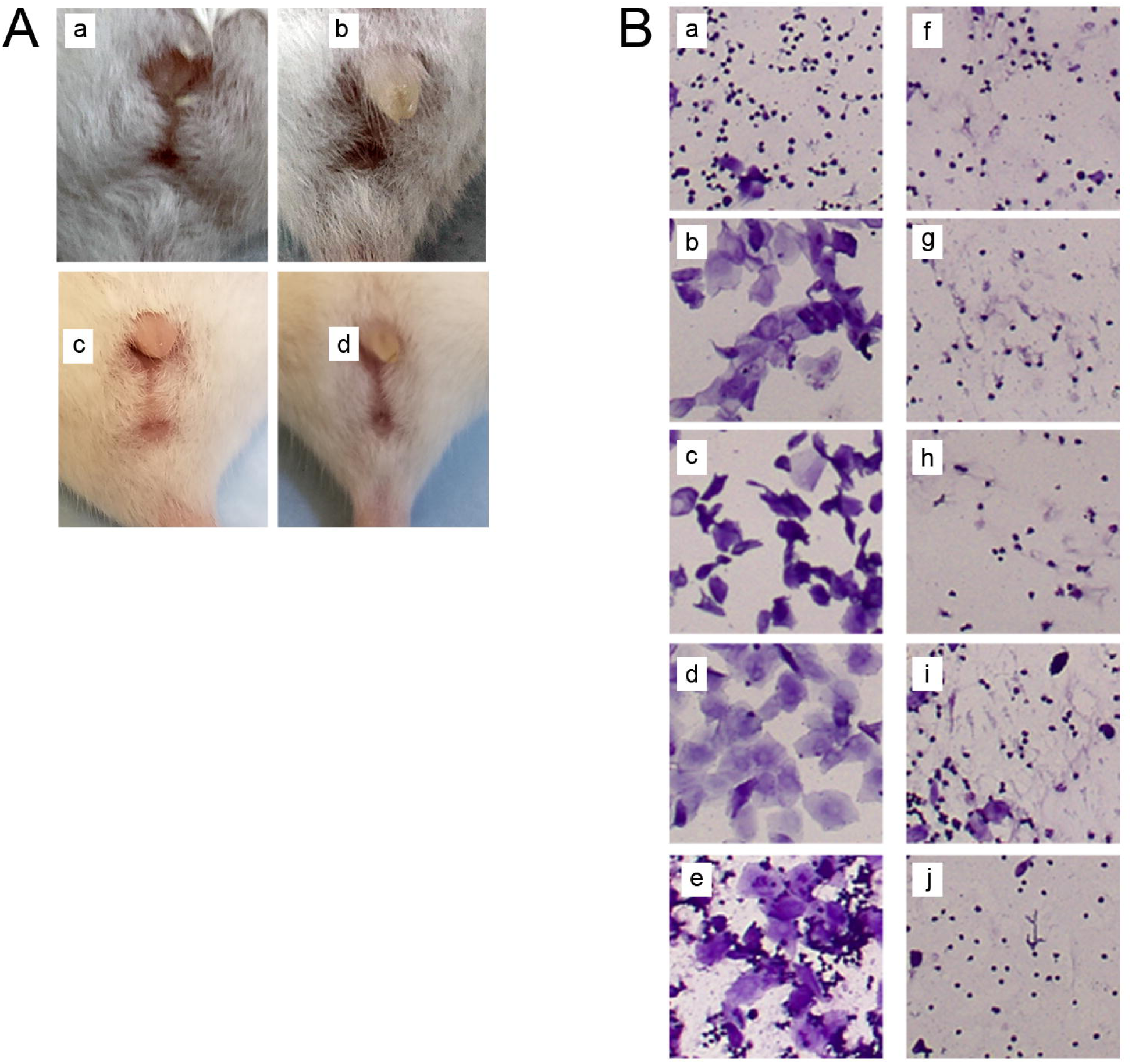
T treatment induces virilization in female mice. A) Clitoromegaly was apparent in the active exposure group (a=control; b=T-treated), but was no longer apparent in the washout group (c=control; d=T-treated). B) Vaginal smears from a cycling, control mouse (a-e) and a T-treated mouse (f-j). Smears were obtained during the fifth week of T injections. Shown here is 5 sequential days of a representative control and T-injected mouse. a=diestrus; b=diestrus into proestrus; c=proestrus; d=estrus; e=metestrus from a control mouse. f-j = T-injected mouse in diestrus.

**Figure 2.**
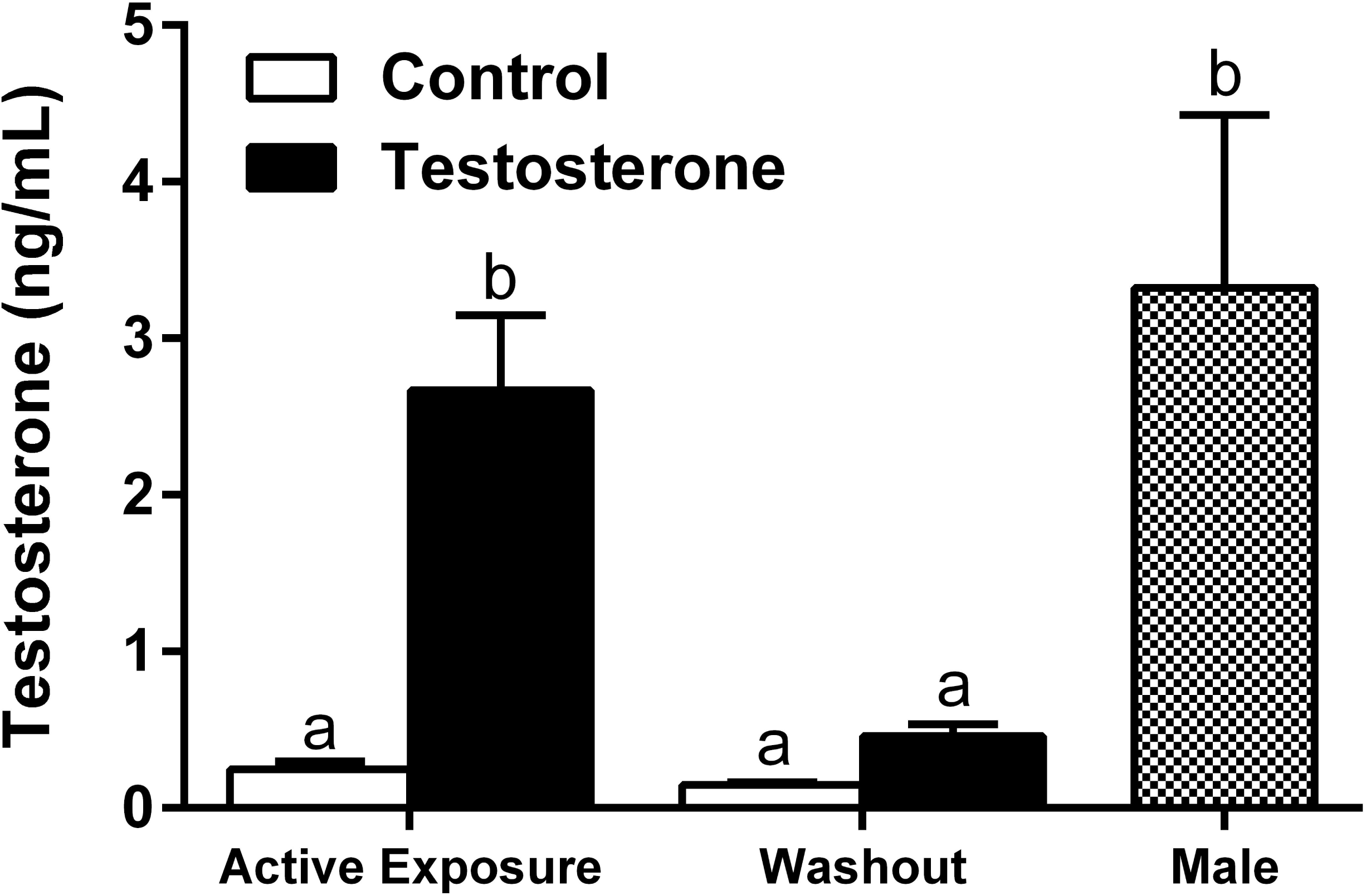
Testosterone cypionate transiently elevates T levels to those of untreated adult males. T levels were significantly higher in the active exposure mice, which were tested following the 6^th^ T injection. T levels declined to baseline levels by 5 weeks after the last injection. Different letters above the bars indicate statistical significance; P<0.0001, as determined by one-way ANOVA.

### Ovaries from T-treated mice are smaller than control ovaries but contain normal complements of follicles and respond to stimulation by gonadotropins

Currently, little is known about the effects of T treatment on the ability of ovaries to respond to gonadotropic stimulation. To investigate this, both the active exposure and washout groups of mice were divided into 3 sub-groups: 1) not stimulated by gonadotropins; 2) stimulated with eCG only; and 3) stimulated with eCG and hCG to induce ovulation. For groups 1 and 2, ovaries were collected after euthanasia, most of the fat and oviducts were removed under a stereoscope, one ovary was fixed and processed for histological analysis, and oocytes were collected from the other ovary. For group 3, ovulated eggs were collected from both oviducts prior to weighing ovaries.

Ovaries from T-treated mice in the active exposure group weighed significantly less than control ovaries whether or not they were stimulated with gonadotropins (Fig. 3A). Notably, the lower ovarian weights were still apparent in the washout group, which more closely mimics the standard of care for T-treated transgender males who wish to obtain functional eggs (Fig. 3B).

**Figure 3.**
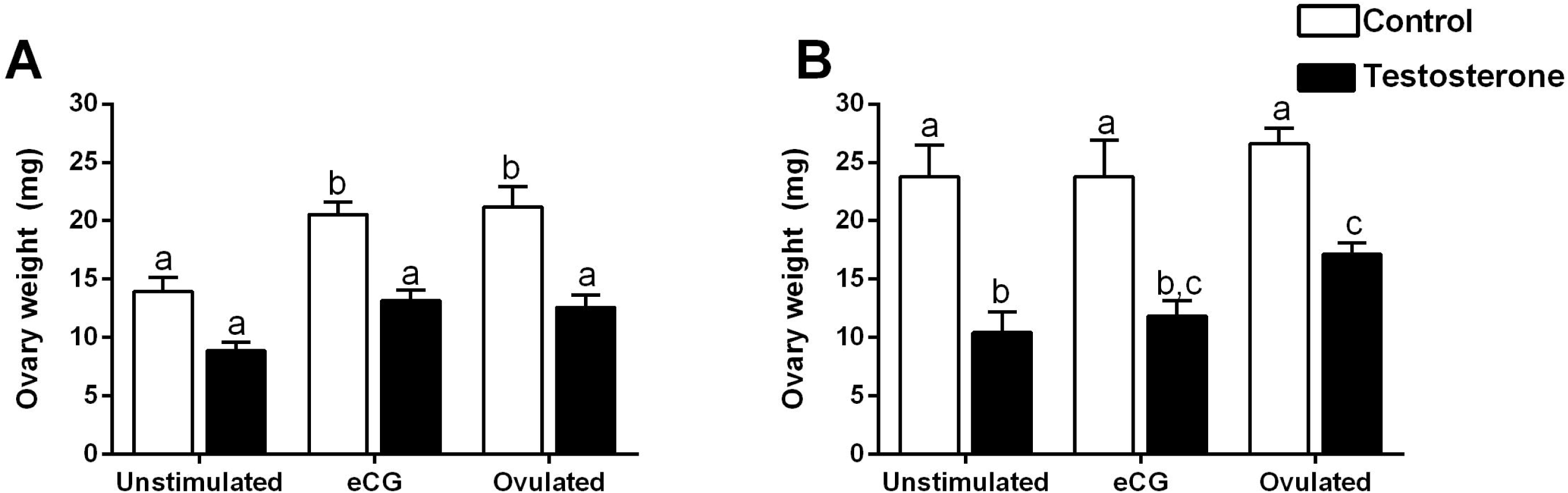
Ovaries from T-treated mice weigh less than control ovaries. A) Ovary weights from mice in the active exposure group. B) Ovary weights from mice in the washout group. Bars with different letters are significantly different (P<0.05), as determined using 2-way ANOVA followed by Bonferroni’s multiple comparison test.

Histological analysis of unstimulated, eCG-stimulated, and ovulated ovaries from the active exposure groups (approximately 12 weeks in age) showed similar follicle morphology and comparable numbers of follicles from T-treated mice with their respective controls, despite the overall smaller size of ovaries in T-treated mice (Fig. 4). Because the numbers of preantral and primary follicles have already been reported to be the same for control and T-treated ovaries (Kinnear et al., 2019), we focused on counting antral follicles of various sizes. We analyzed follicles in detail from 3 mice per group, one ovary from each mouse. In the unstimulated group, T-treated and control ovaries contained similar numbers of antral follicles, as well as similar numbers of atretic follicles. Most of the atretic follicles were in the 250-320 μm size range, while there were almost no atretic follicles in the preovulatory size range. The major difference between T-treated, unstimulated ovaries and control ovaries was a significantly lower number of corpora lutea (CLs) in the T-treated group compared with controls (Fig. 4). The eCG-stimulated ovaries contained significantly more preovulatory follicles than the unstimulated ovaries in both control and T-treated mice and contained 0-1 atretic preovulatory follicles. Similar to unstimulated ovaries, ovaries from T-treated mice contained fewer CLs than controls. Most of the CLs present in the T-treated ovaries were likely to be from cycles that occurred prior to T treatment, as they were eosinophilic rather than basophilic (Gaytan et al., 2017) and, in general, located deep within the ovary rather than at the periphery (Fig. 4). The one exception was a T-treated mouse that ovulated in response to the eCG injection; this mouse contained mostly basophilic CLs (Fig. 4A). We did not evaluate ovulated ovaries in detail with histology.

**Figure 4.**
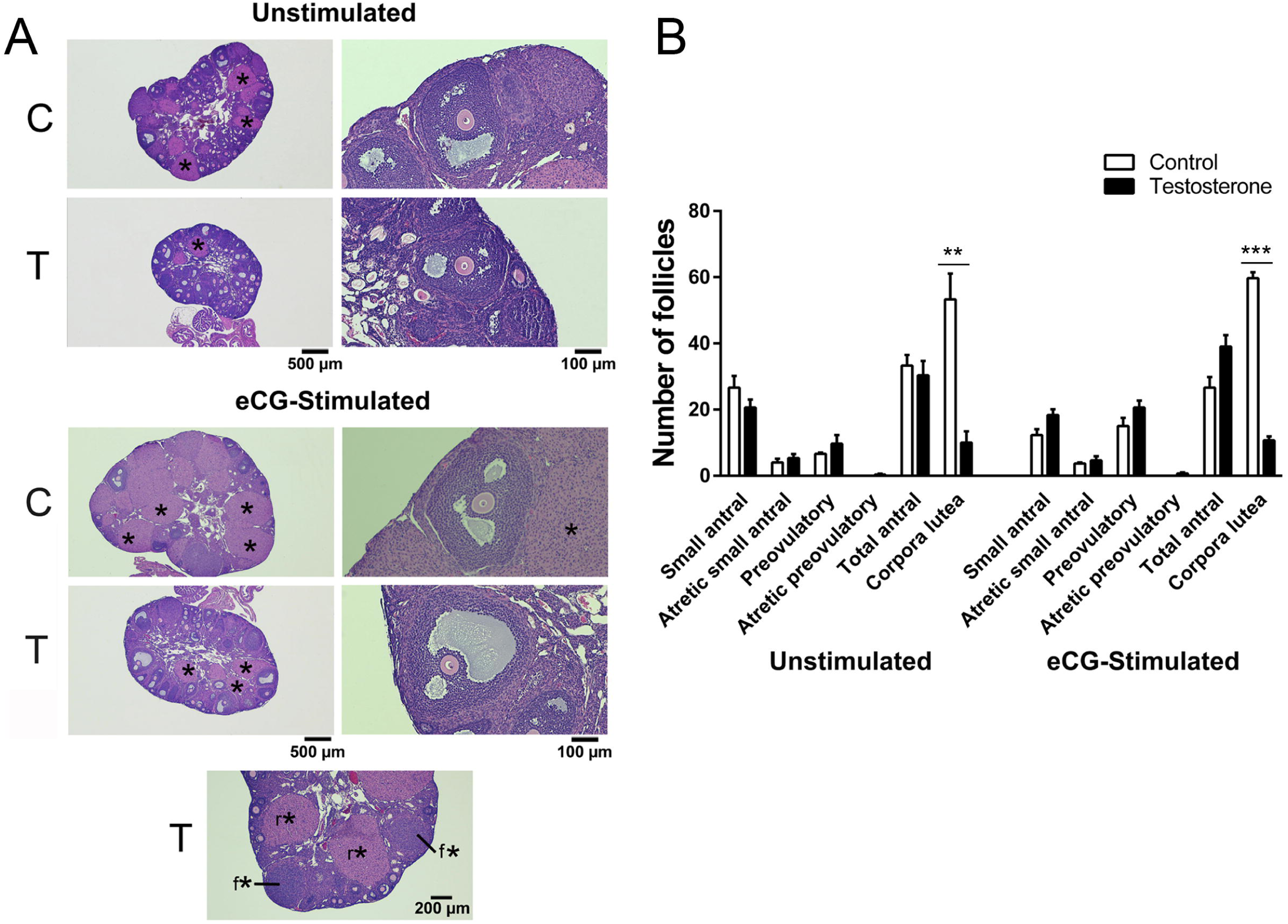
Ovaries from T-treated mice have normal complements of antral follicles but fewer corpora lutea. A) Histology sections showing representative images through ovaries from unstimulated and eCG-stimulated control and T-treated mice from the active exposure group using 2X, 4X, or 10X objectives. C=control; T=testosterone; * = corpus luteum. The bottommost image is from a T-treated mouse that ovulated in response to eCG stimulation, showing fresh CLs (f*) and residual CLs (r*). B) Quantification of antral follicles in unstimulated and eCG-stimulated control and T-treated mice. “Small antral” follicles measured ∼250-320 μm in diameter; “preovulatory” follicles measured >320 μm in diameter. ** P<0.01 (unpaired t-test); *** P<0.0001 (unpaired t-test). The increase in preovulatory follicles from eCG-stimulated ovaries compared to unstimulated ovaries is significant (P<0.05; 2-way ANOVA followed by Bonferroni’s multiple comparison test). Data are mean ± SEM.

Measurements of estradiol-17β (E2) levels from trunk blood obtained at the time of sacrifice provided further evidence that T-stimulated ovaries responded to gonadotropins: E2 was low in the unstimulated groups, increased in response to eCG injection, then fell back to basal levels after ovulation (Fig. 5). E2 levels in T-treated mice in both the active exposure group and the washout group were lower than respective controls in response to eCG stimulation, and E2 levels in the eCG-stimulated mice from the washout group were ∼3X higher than in the active exposure group (Fig. 5), though the basal levels in the unstimulated and ovulated groups were similar to those in the active exposure group.

**Figure 5.**
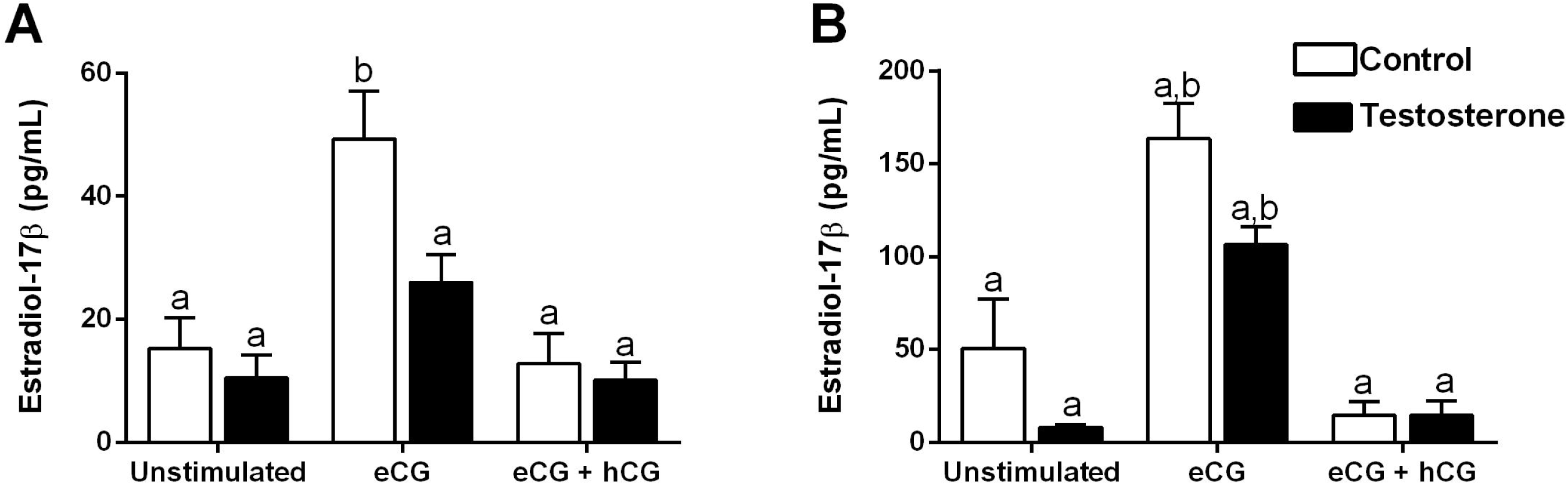
eCG stimulates E2 production in control and T-treated mice in both the active exposure and washout groups. Bars with different letters are significantly different (P<0.05; 2-way ANOVA, Bonferroni’s multiple comparison post-test). Data are mean ± SEM.

### T-treated mice contain comparable numbers of meiotically competent oocytes and ovulate similar numbers of fertilizable, mature eggs as controls

To examine if the follicles from T-treated mice contain normal oocytes, we first collected immature oocytes from ovaries of unstimulated and eCG-stimulated mice, and then collected ovulated eggs from the groups that were injected with both eCG and hCG. In the active exposure group, we recovered significantly more immature oocytes from the T-exposed ovaries than from their respective controls (Fig. 6A). In the washout group, we obtained similar numbers of immature oocytes from T-treated and control ovaries, though statistical analysis was not possible in this group due to the small sample size (n=2; Fig. 6B). Both active exposure and washout groups contained similar numbers of ovulated eggs for T-treated and control mice (Fig. 6A,B). Immature oocytes from T-treated ovaries (active exposure and washout groups) underwent germinal vesicle breakdown (GVBD), extruded first polar bodies, and formed morphologically normal meiotic spindles in culture (Fig. 6C,D). The proportion of oocytes with intact spindle structure vs poor spindle structure - characterized by degeneration or misaligned chromosomes - was similar between the groups (Fig. 6D). The diameters of in vitro matured eggs from the active exposure T group were the same as controls (Fig. 6E).

**Figure 6.**
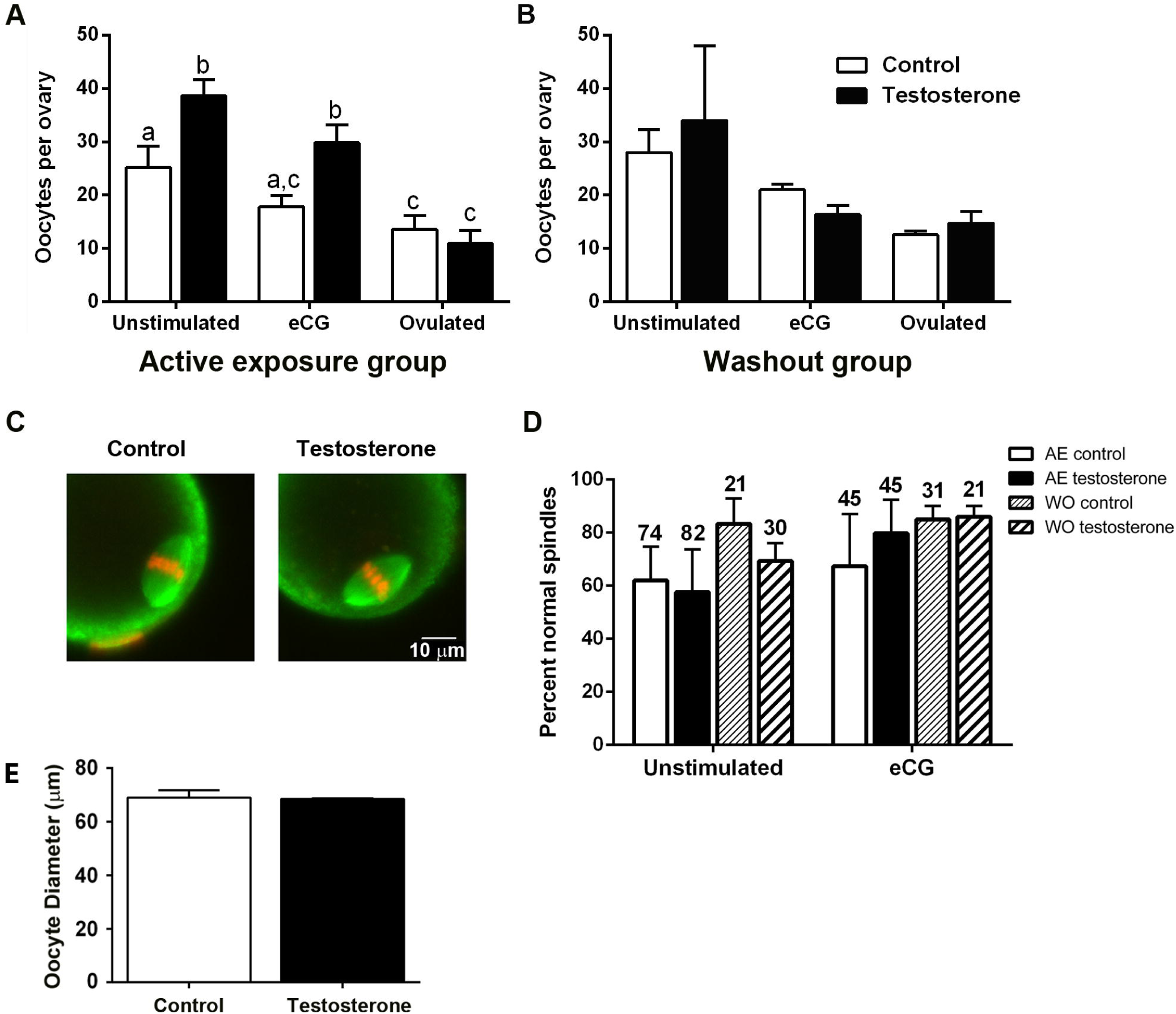
T-treated mice produce meiotically competent oocytes and ovulate comparable numbers of eggs as controls. A,B) Numbers of oocytes and ovulated eggs recovered per ovary in the active exposure (A) and washout group (B). Bars are mean ± SEM; P<0.05 was considered significant, as determined by two-way ANOVA followed by Bonferroni’s multiple comparison test. C) Representative meiotic spindles from in vitro matured, eCG-stimulated ovaries from the active exposure group. Green = tubulin; Red = DNA. D) Percentage of eggs that formed normal meiotic spindles following in vitro maturation. Bars are mean ± SEM. Numbers over each bar are the total number of in vitro matured eggs. AE = active exposure; WO = washout. E) Diameters of in vitro matured eggs in control vs. T-treated mice.

We tested the fertilizability of ovulated eggs using in vitro fertilization by evaluating the number of 2-cell embryos that were observed 24 hrs after insemination. Overall, cleavage to the 2-cell stage was comparable between T-treated and control mice, and was similar to percentages of fertilized eggs obtained from the washout group (Fig. 7).

**Figure 7.**
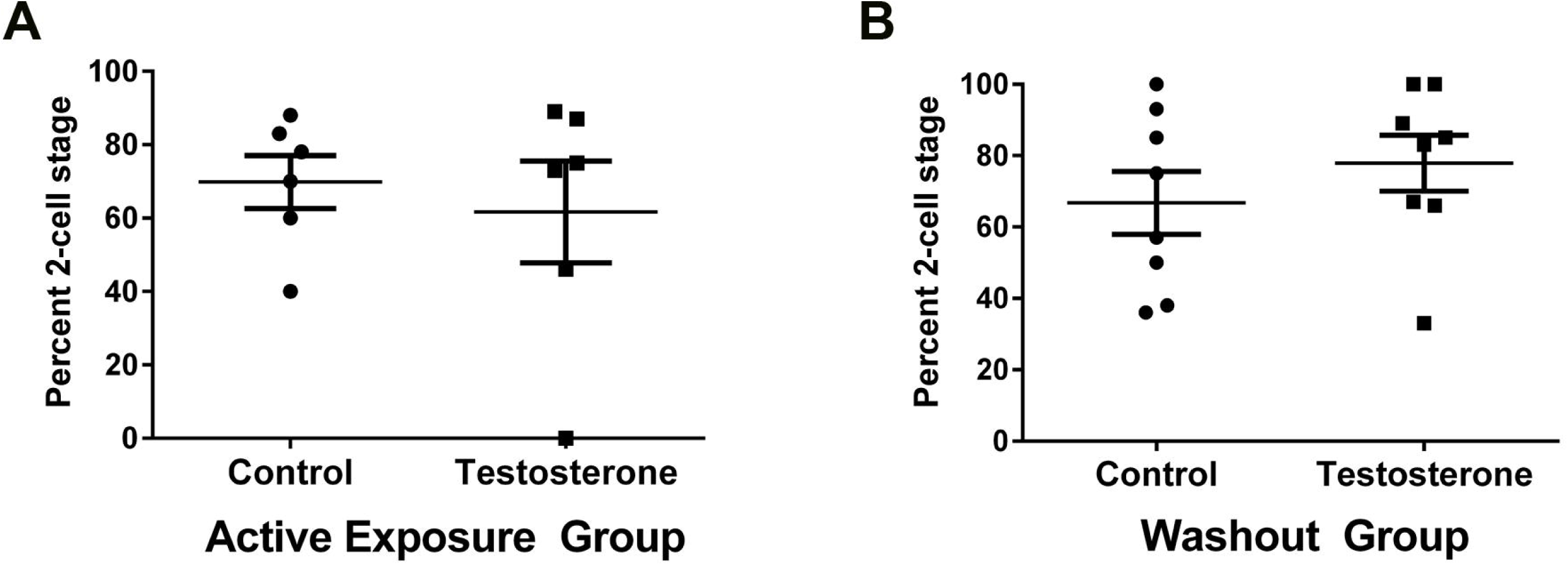
Eggs from T-treated mice are fertilizable. Completed fertilization rates, defined as the number of 2-cell embryos per the number of inseminated eggs, in control vs. T-treated mice in both active exposure (A) and washout (B) groups. Each dot represents a single mouse. Horizontal bars are the mean and the vertical bars are ± SEM.

## Discussion

Transgender men who have been undergoing testosterone therapy and who wish to obtain eggs for fertilization or freezing are generally stimulated with gonadotropins. This is usually done following a period of T cessation that is sufficient for the menstrual cycle to resume (Adeleye et al., 2019; Broughton and Omurtag, 2017; Leung et al., 2018). There is strong evidence that transgender men who have taken T can produce viable, developmentally competent eggs after discontinuing its use (Adeleye et al., 2019; Broughton and Omurtag, 2017; Leung et al., 2018; Light et al., 2014), but to date, there are no publications regarding the quality and developmental capacity of eggs retrieved from transgender men who remain on HT. Here, we show that treating female mice with T cypionate weekly for 6 weeks does not impair the fertilizability of their eggs, and our results suggest that T treatment does not need to stop before gonadotropic stimulation.

T cypionate elevated blood T levels to those found in male mice, stopped the estrous cycle, and caused significant clitoral growth, changes that are commonly observed in transgender males on HT (Unger, 2016). It was noteworthy that the clitoromegaly did not persist in the washout group, which was surprising because it is generally recognized that this is a permanent change in human females exposed to high levels of T (Cabrera and Rogol, 2013; Sielert et al., 2013). It is possible that the regression of clitoromegaly upon T withdrawal in our mice represents a difference between mice and humans; however, careful measurements using a larger sample size would need to be done to determine this definitively.

T-treated mice had smaller ovaries than controls, which is likely due to the greatly reduced number of CLs in these mice. A recent study reported a complete absence of CLs in T-treated female mice (Kinnear et al., 2019). The finding that our T-treated mice had CLs at all was unexpected, as we used a concentration of T cypionate that was similar to the mid-range effective dose used by Kinnear et al. In general, the eosinophilic staining of CLs we observed in T-treated mice was consistent with residual rather than freshly ovulated CLs (Gaytan et al., 2017) and there were considerably fewer CLs in our T-treated mice than in controls, suggesting that the estrous cycle was indeed inhibited in our mice, as was also shown by vaginal cytology. Smaller ovaries were also apparent in T-treated mice in the washout group, which likewise contained significantly fewer CLs than controls, suggesting that these mice had only recently begun cycling prior to ovary harvest and therefore had many fewer ovulations than controls.

Although they weighed less than controls, the ovaries from T-treated mice contained a complement of histologically normal antral follicles that were similar to controls. Unlike the study by Kinnear et al. (2019), who reported a higher incidence of late-stage atretic, cyst-like follicles, we only observed a single preovulatory atretic follicle in 3/6 ovaries examined, which was not significantly different from controls. Rather, the majority of atretic antral follicles we observed came from follicles that were in the ∼250-320 μm diameter size range, not yet preovulatory, and the percentages were not different between control and T-treated ovaries. T-treated ovaries that were stimulated by eCG had more preovulatory follicles than unstimulated ovaries, and these numbers were comparable to the number of preovulatory follicles observed in control mice. One of the T-treated mice unexpectedly ovulated in response to eCG stimulation. There is evidence that T induces the expression of FSH receptors in granulosa cells (Garcia-Velasco et al., 2012; Liu et al., 2015; Sen et al., 2014), and if this occurred in our mice, then it is possible that the T-treated follicles were sensitized to eCG such that spontaneous ovulation occurred prior to the administration of hCG.

T-treated mice responded to eCG treatment by producing E2. Interestingly, T-treated mice in the active exposure group did not elevate E2 to the same extent as controls. The difference in E2 levels after eCG was surprising because a similar number of oocytes were collected after stimulation. In human IVF cycles, the peak E2 level achieved directly correlates with the number of oocytes collected at the time of retrieval (Chenette et al., 1990), and during reported IVF cycles in transgender males, the peak E2 levels, as well as the number of oocytes collected, have been reported to be similar to female controls, although higher doses of gonadotropins were used for transgender males (Leung et al., 2018). Lower E2 levels here may reflect the lower number of CLs in the T group. Though it is thought that CLs are rapidly inactivated after each cycle in mice (Accialini et al., 2017), it is possible that the residual CLs retain some ability to produce E2 in response to eCG.

Ovaries from T-treated mice produced meiotically competent oocytes. Oocytes retrieved from unstimulated and eCG-stimulated, T-treated ovaries were able to mature to the metaphase II stage in culture, forming morphologically normal meiotic spindles. This is consistent with a descriptive study of human oocytes collected from the ovarian cortex of HT-exposed transgender males, in which oocyte meiotic spindle structure after in vitro maturation was found to be normal (Lierman et al., 2017). In addition, T-treated mice in both the active exposure and washout groups ovulated similar numbers of eggs in response to eCG and hCG injection as controls. These eggs fertilized to the same extent as controls and cleaved to the 2-cell stage. One exception in the active exposure group was a single T-treated mouse that only produced 3 poor-quality eggs, none of which fertilized. One hypothesis is that this mouse ovulated prematurely, as there was some histological evidence in a different T-treated mouse of premature ovulation after eCG. There was insufficient data to fully explore this isolated scenario, and the other T-treated mice produced fertilizable eggs.

In conclusion, we provide evidence showing that female mice produce normal, fertilizable eggs after testosterone exposure, whether T levels are low after a washout period or high during active exposure. One limitation in our study could be that we only exposed mice to T for 6 weeks. While this length of exposure was sufficient to produce phenotypes characteristic of human transgender males exposed to T, it may not completely mimic the human situation, in which many transgender males seeking fertility treatment have been on HT for several years. Further studies that expose mice to T for longer periods of time would help confirm that its effects are not detrimental to the reproductive process. Despite this concern, our results provide promising data that could help influence the treatment options for transgender men seeking fertility treatment. If testosterone has no detrimental impact on ovarian function, transgender males will have greater flexibility in making reproductive decisions. It is important to note, however, that if a transgender male plans to become pregnant by spontaneous pregnancy or use of assisted reproductive technology, testosterone must be discontinued due to its teratogenic effects (De Roo et al., 2016). Our study suggests that the current practice of T cessation prior to ovarian stimulation and surgical oocyte retrieval may not be necessary when the transgender male does not plan to carry the pregnancy at that time, and could potentially help serve as the basis for human trials to examine this current clinical practice.

## Authors’ Roles

C.B.B. designed the study, acquired and interpreted the data, and helped draft the article. T.F.U. contributed to the study design, data acquisition, and article review. L.L. contributed to data acquisition and article review. L.M.M. designed the study, contributed to data acquisition and interpretation, and helped draft the article.

## Acknowledgements

The authors thank Dr. Daniel Grow, M.D. and director of the Center for Advanced Reproductive Services Fellowship Program at UConn Health; and Drs. Laurinda Jaffe, Ph.D., Bruce White, Ph. D, and John Nulsen, M.D. for support and helpful advice, discussions, and suggestions on the manuscript.

## References

Accialini P, Hernandez S, Abramovich D, Tesone M. The Rodent Corpus Luteum. In Meidan R (ed) The Life Cycle of the Corpus Luteum. 2017 Springer International Publishing, New York, NY, pp 117-131.

Adeleye AJ, Cedars MI, Smith J, Mok-Lin E. Ovarian stimulation for fertility preservation or family building in a cohort of transgender men. J Assist Reprod Genet 2019;36: 2155–2161.

Armuand G, Dhejne C, Olofsson JI, Rodriguez-Wallberg KA. Transgender men’s experiences of fertility preservation: a qualitative study. Hum Reprod 2017;32: 383–390.

Broughton D, Omurtag K. Care of the transgender or gender-nonconforming patient undergoing in vitro fertilization. Int. J Transgenderism 2017;18: 372–375.

Cabrera SM, Rogol AD. Testosterone exposure in childhood: discerning pathology from physiology. Expert Opin Drug Saf 2013;12: 375–388.

Chenette PE, Sauer MV, Paulson RJ. Very high serum estradiol levels are not detrimental to clinical outcome of in vitro fertilization. Fertil Steril 1990;54: 858–863.

De Roo C, Tilleman K, T’Sjoen G, De Sutter P. Fertility options in transgender people. Int Rev Psych 2016;28: 112–119.

De Roo C, Tilleman K, Vercruysse C, Declercq H, T’Sjoen G, Weyers S, De Sutter P. Texture profile analysis reveals a stiffer ovarian cortex after testosterone therapy: a pilot study. J Assist Reprod Genet 2019;36: 1837–1843.

Flores AR, Herman, J.L., Gates, G.J., Brown, T.N.T. How many adults identify as transgender in the United States? Los Angeles, CA: The Williams Institute 2016.

Garcia-Velasco JA, Rodriguez S, Agudo D, Pacheco A, Schneider J, Pellicer A. FSH receptor in vitro modulation by testosterone and hCG in human luteinized granulosa cells. Eur J Obstet Gynecol Reprod Biol 2012;165: 259–264.

Gaytan F, Morales C, Leon S, Heras V, Barroso A, Avendano MS, Vazquez MJ, Castellano JM, Roa J, Tena-Sempere M. Development and validation of a method for precise dating of female puberty in laboratory rodents: The puberty ovarian maturation score (Pub-Score). Sci Rep 2017;7: 46381.

Goldman JM, Murr AS, Cooper RL. The rodent estrous cycle: characterization of vaginal cytology and its utility in toxicological studies. Birth Defects Res B Dev Reprod Toxicol 2007;80: 84–97.

Hembree WC, Cohen-Kettenis PT, Gooren L, Hannema SE, Meyer WJ, Murad MH, Rosenthal SM, Safer JD, Tangpricha V, T’Sjoen GG. Endocrine Treatment of Gender-Dysphoric/Gender-Incongruent Persons: An Endocrine Society Clinical Practice Guideline. J Clin Endocrinol Metab 2017;102: 3869–3903.

Ikeda K, Baba T, Noguchi H, Nagasawa K, Endo T, Kiya T, Saito T. Excessive androgen exposure in female-to-male transsexual persons of reproductive age induces hyperplasia of the ovarian cortex and stroma but not polycystic ovary morphology. Hum Reprod 2013;28: 453–461.

Insogna IG, Ginsburg E, Srouji S. Fertility Preservation for Adolescent Transgender Male Patients: A Case Series. J Adolesc Health 2020.

Kinnear HM, Constance ES, David A, Marsh EE, Padmanabhan V, Shikanov A, Moravek MB. A mouse model to investigate the impact of testosterone therapy on reproduction in transgender men. Hum Reprod 2019:34: 2009–2017.

Leung A, Sakkas D, Pang S, Thornton K, Resetkova N. ART outcomes in female to male transgender patients: a new frontier in reproductive medicine. Fert Steril 2018;109: e35.

Lierman S, Tilleman K, Braeckmans K, Peynshaert K, Weyers S, T’Sjoen G, De Sutter P. Fertility preservation for trans men: frozen-thawed in vitro matured oocytes collected at the time of ovarian tissue processing exhibit normal meiotic spindles. J Assist Reprod Genet 2017;34: 1449–1456.

Light AD, Obedin-Maliver J, Sevelius JM, Kerns JL. Transgender men who experienced pregnancy after female-to-male gender transitioning. Obstet Gynecol 2014;124: 1120–1127.

Liu T, Cui YQ, Zhao H, Liu HB, Zhao SD, Gao Y, Mu XL, Gao F, Chen ZJ. High levels of testosterone inhibit ovarian follicle development by repressing the FSH signaling pathway. J Huazhong Univ Sci Technolog Med Sci 2015;35: 723–729.

Luthy K, Williams C, Freeborn D, Cook A. Comparison of testosterone replacement therapy medications in the treatment of hypogonadism. J Nurse Pract 2017;13: 241–249.

Mehlmann LM, Kline D. Regulation of intracellular calcium in the mouse egg: calcium release in response to sperm or inositol trisphosphate is enhanced after meiotic maturation. Biol Reprod 1994;51: 1088–1098.

Mehlmann LM, Uliasz T, Lowther KM. SNAP23 is required for constitutive and regulated exocytosis in mouse oocytes. Biol Reprod 2019;101: 338–346.

Meyer WJ. World professional association for transgender health’s standards of care requirements of hormone therapy for adults with gender identity disorder. Int J Transgenderism 2009;11: 127–132.

Moravek MB. Fertility preservation options for transgender and gender-nonconforming individuals. Curr Opin Obstet Gynecol 2019;31: 170–176.

Moravek MB, Kinnear HM, George J, Batchelor J, Shikanov A, Padmanabhan V, Randolph JF. Impact of Exogenous Testosterone on Reproduction in Transgender Men. Endocrinology 2020;161.

Nahata L, Chen D, Moravek MB, Quinn GP, Sutter ME, Taylor J, Tishelman AC, Gomez-Lobo V. Understudied and Under-Reported: Fertility Issues in Transgender Youth-A Narrative Review. J Pediatr 2019;205: 265–271.

Neblett MF, 2nd, Hipp HS. Fertility Considerations in Transgender Persons. Endocrinol Metab Clin North Am 2019;48: 391–402.

Quinn VP, Nash R, Hunkeler E, Contreras R, Cromwell L, Becerra-Culqui TA, Getahun D, Giammattei S, Lash TL, Millman A et al. Cohort profile: Study of Transition, Outcomes and Gender (STRONG) to assess health status of transgender people. BMJ Open 2017;7: e018121.

Rodriguez-Wallberg KA, Dhejne C, Stefenson M, Degerblad M, Olofsson JI. Preserving eggs for men’s fertility. a pilot experience with fertility preservation for female-to-male transsexuals in Sweden. Fert Steril 2014;102: e160–e161.

Sen A, Prizant H, Light A, Biswas A, Hayes E, Lee HJ, Barad D, Gleicher N, Hammes SR. Androgens regulate ovarian follicular development by increasing follicle stimulating hormone receptor and microRNA-125b expression. Proc Natl Acad Sci U S A 2014;111: 3008–3013.

Sielert L, Liu C, Nagarathinam R, Craig LB. Androgen-producing steroid cell ovarian tumor in a young woman and subsequent spontaneous pregnancy. J Assist Reprod Genet 2013;30: 1157–1160.

Unger CA. Hormone therapy for transgender patients. Transl Androl Urol 2016;5: 877–884.

Van Den Broecke R, Van Der Elst J, Liu J, Hovatta O, Dhont M. The female-to-male transsexual patient: a source of human ovarian cortical tissue for experimental use. Hum Reprod 2001;16: 145–147.

Wierckx K, Van Caenegem E, Pennings G, Elaut E, Dedecker D, Van de Peer F, Weyers S, De Sutter P, T’Sjoen G. Reproductive wish in transsexual men. Hum Reprod 2012;27: 483–487.

